# Casein kinase 1D encodes a novel drug target in Hedgehog-GLI driven cancers and tumor-initiating cells resistant to SMO inhibition

**DOI:** 10.1101/2021.06.17.448773

**Authors:** Elisabeth Peer, Sophie Karoline Aichberger, Filip Vilotic, Wolfgang Gruber, Thomas Parigger, Sandra Grund-Gröschke, Dominik P. Elmer, Florian Rathje, Andrea Ramspacher, Mirko Zaja, Susanne Michel, Svetlana Hamm, Fritz Aberger

**Author notes:** present address: EVER Pharma GmbH, Unterach am Attersee, Austria. present address: Immatics Biotechnologies GmbH, Tübingen, Germany (Sandra Grund-Gröschke).

## Abstract

**Background:** Aberrant activation of the Hedgehog (HH)/GLI pathway in stem-like tumor initiating cells (TIC) is a frequent oncogenic driver signal in various human malignancies. Remarkable efficacy of anti-HH therapeutics led to the approval of HH inhibitors targeting the key pathway effector Smoothened (SMO) in basal cell carcinoma and acute myeloid leukemia. However, frequent development of drug resistance and severe adverse effects of SMO inhibitors pose major challenges that require alternative treatment strategies targeting HH/GLI in TIC downstream of SMO. We therefore investigated members of the casein kinase 1 (CSNK1) family as novel drug targets in HH/GLI driven malignancies.

**Methods:** We genetically and pharmacologically inhibited CSNK1D in HH-dependent cancer cells displaying either sensitivity or resistance to SMO inhibitors. To address the role of CSNK1D in oncogenic HH signaling and tumor growth and initiation, we quantitatively analyzed HH target gene expression, performed genetic and chemical perturbations of CSNK1D activity and monitored oncogenic transformation of TIC *in vitro* and *in vivo* using 3D clonogenic tumor spheroid assays and xenograft models.

**Results:** We show that CSNK1D plays a critical role in controlling oncogenic GLI activity downstream of SMO. We provide evidence that inhibition of CSNK1D interferes with oncogenic HH signaling in both SMO-inhibitor sensitive and resistant tumor settings. Furthermore, genetic and pharmacologic perturbation of CSNK1D decreases the clonogenic growth of GLI-dependent tumor-initiating cancer cells *in vitro* and *in vivo*.

**Conclusions:** Pharmacologic targeting of CSNK1D represents a novel therapeutic approach for the treatment of both SMO inhibitor sensitive and resistant tumors.

## 1. Introduction

Cancer tissues typically display a hierarchical organization reflected by the existence of rare yet highly malignant stem-like tumor initiating cells (TIC) and more abundant differentiated progeny. TICs are key to cancer initiation and growth, display less sensitivity to chemotherapeutics and are endowed with self-renewal and metastatic capacity. They are frequently enriched in advanced, aggressive and/or resistant tumors, and cells isolated from distant metastases often show a TIC phenotype [1,2]. A better understanding of the key molecular drivers and pathways accounting for the malignant properties of TICs is, therefore, of high therapeutic relevance.

On a molecular level, the Hedgehog (HH)/GLI signaling pathway has been crucially implicated in the regulation of self-renewal, disseminating and tumor initiating capacity of TICs [3–7]. This pivotal role of HH/GLI in TICs makes targeted pharmacological inhibition of HH/GLI signaling a promising therapeutic strategy to combat some of the major challenges in oncology such as patients′ relapse and metastases formation.

The Hedgehog (HH) signaling pathway is of central importance during embryonic development and regeneration, where it controls a multitude of biological processes such as cell proliferation, differentiation, survival, cell metabolism and stem cell fate. In line with its various key roles, aberrant activity of the pathway in humans is causally linked with several developmental syndromes and malignancies [8–11]. Canonical Hedgehog signaling starts with binding of the HH ligand protein to its receptor Patched (PTCH), a twelve-transmembrane domain protein, which represses HH signaling in its unliganded state by inhibiting the transport of the G-protein-coupled receptor-like transmembrane protein Smoothened (SMO) into the primary cilium [12]. In its activated state, SMO promotes the formation of active glioma-associated oncogene homolog (GLI) transcription factors by releasing GLI2/3 from their repressor Suppressor of Fused (SUFU), thereby preventing proteolytic processing of GLI2/3. The unprocessed, full-length GLI proteins are then able to activate expression of HH target genes, which promote tumor formation by inducing proliferation, anti-apoptotic signals, metastasis, cancer stem cell (CSC) self-renewal as well as expression of GLI1, leading to a strong positive feedback circuit. Furthermore, GLI1 expression serves as a reliable readout for HH/GLI pathway activation (for reviews, see [13–17]).

The first inhibitors of oncogenic HH/GLI that were approved by the FDA for the treatment of advanced and metastatic basal cell carcinoma (BCC) were the small molecule SMO inhibitors (SMOi) vismodegib and sonidegib [18–22]. More recently, the third SMOi glasdegib has been approved for the treatment of acute myeloid leukemia in combination with low-dose cytarabine [23]. Although the SMO inhibitors demonstrate remarkable therapeutic efficacy, they are also associated with severe side effects such as muscle cramps, weight and hair loss and taste disturbance. These adverse effects very frequently force discontinuation of the treatment. However, drug withdrawal is often followed by tumor relapse, due to the existence of resistant tumors cells. Here, it is noteworthy that SMO-inhibitors alone rarely eliminate all tumor cells, which allows residual tumor cells to persist and regrow [24]. A substantial proportion of BCC patients showing SMOi resistance express mutant SMO variants. Such mutations in SMO occur either directly in the SMO inhibitor-binding pocket or outside of the binding pocket in pivotal regions of the transmembrane-helices, abolishing or attenuating the antagonistic activity of SMOi drugs. Additionally, resistance to SMO inhibitors can be caused by GLI2 amplification or genetic loss of the GLI repressor SUFU [25–29]. Further, the antagonistic activity of SMO inhibitors can be bypassed by signaling shifts towards phosphatidyl inositol-3 kinase (PI3K), activation of mitogen activated protein kinase (MAPK) signaling or by induction of atypical protein kinase C (PKC) activity, the latter of which enhancing transcriptional GLI activity independent of SMO function [28,30–33]. Along the same line, Whitson *et al*. demonstrated that serum response factor (SRF) and the coactivator megakaryoblastic leukemia 1 (MLK1) together with GLI1 activation drives non-canonical Hedgehog signaling and induces SMOi resistance in BCC patients [31].

Considering the numerous kinases and signaling pathways, also including EGF-MEK-ERK, FGF, PDGF, TGF, Wnt, JAK/STAT, or DYRK [34–43] that interact with the HH signaling pathway and are often integrated and translated into synergistically regulated output signals via the GLI transcription factors, we focused in this study on members of the casein kinase family with the goal of identifying novel druggable target proteins to inhibit oncogenic GLI function. Members of the casein kinase (CSNK) family have been shown to play multiple and diverse roles in HH signaling [44]. For instance, CSNK1 alpha (CSNK1A) negatively regulates insect and vertebrate HH signaling by phosphorylating GLI3, and thereby inducing proteolytic processing of GLI3 full-length protein into C-terminally truncated repressor forms [45,46]. By contrast, CSNK1A has also been shown to phosphorylate SMO and thereby contribute to activation of HH signaling [47]. Similarly, CSNK2 has been identified as driver signal and therapeutic target in HH-dependent medulloblastoma [48]. Further evidence for an activating role of CSNK proteins came from studies in in *Drosophila* showing that fly homologues of CSNK1 can phosphorylate and thereby stabilize the GLI homologue Ci, increasing the transcriptional output of hh signaling in insects [49,50]. Together, these data suggest that pharmacologic inhibition of selected CSNK family members may be a promising therapeutic approach for HH-GLI driven cancers, particularly malignancies with SMOi resistance.

Here, we identify Casein Kinase 1-Delta (CSNK1D) as positive regulator of oncogenic GLI function and show that pharmacological as well as genetic targeting of CSNK1D is sufficient to abrogate HH/GLI signaling in both SMOi-sensitive and SMOi-resistant cancer cells driven by oncogenic GLI. Furthermore, we demonstrate a requirement of CSNK1D for GLI-dependent TIC properties *in vitro* and *in vivo* and introduce a novel CSNK1D inhibitor with therapeutic activity against GLI-driven cancer cells.

## 2. Results

### 2.1. Genetic perturbation of CSNK1D interferes with canonical, oncogenic HH/GLI signaling in medulloblastoma cells

To analyze whether CSNK1D is involved in the regulation of oncogenic HH signaling, we performed an shRNA-mediated knockdown of CSNK1D in a HH responsive human medulloblastoma cell line (Daoy). In Daoy cells canonical HH signaling can be activated and inhibited by treatment with a small molecule Smoothened agonist (SAG) and SMO antagonists, respectively [51,52] (Fig 1A). GLI1 protein as well as mRNA levels of the HH target genes GLI1 and HHIP whose expression significantly increases upon the addition of SAG were used as readout parameter for HH pathway activity. In addition, Daoy cells express high levels of CSNK1D, which can be inhibited via shRNA-mediated knockdown (Fig 1B). Of note, knockdown of CSNK1D was sufficient to reduce HH target gene and GLI1 protein expression in Daoy medulloblastoma cells, suggesting that CSNK1D is required for canonical HH signaling and HH target gene activation (Fig 1B and 1C). Knockdown of CSNK1D showed no effect on primary cilia, suggesting that loss of CSNK1D directly interferes with GLI activation rather than via interfering with the formation of primary cilia, which serve as essential organelles in the reception and transduction of HH signaling upstream of GLI proteins (suppl. Fig. S1A, B) [53].

**Figure 1.**
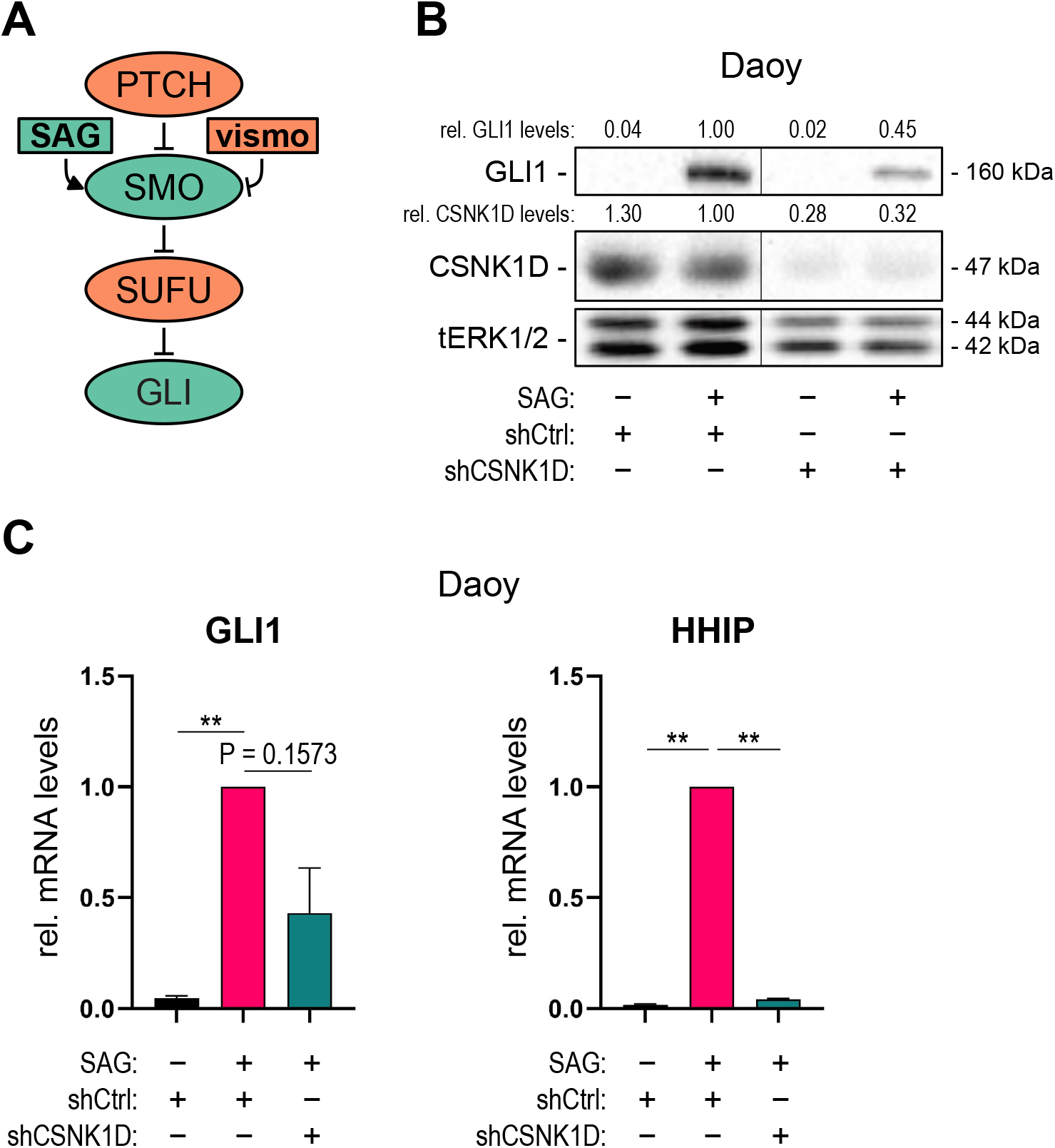
RNAi-mediated inhibition of CSNK1D interferes with canonical oncogenic HH/GLI signaling. **(A)** Simplified schematic illustration of the canonical HH/GLI signaling pathway with pathway activators in green and inhibitors in red. PTCH and SUFU inactivate the pathway, while it is turned on by the central pathway activator SMO causing activation of the GLI transcription factor. Pharmacologically, the SMO agonist SAG activates the pathway, while it is inhibited by the SMO inhibitor vismodegib. **(B)** Representative Western blot analysis of GLI1 and CSNK1D in Daoy medulloblastoma cells. Cells were lentivirally transduced with shCSNK1D or control shRNA (shCtrl) and treated with or without SAG [100 nM]. Relative quantification of Western blot bands was conducted via densitometric image analysis using Image Lab 5.0 software (Bio-Rad, Vienna, Austria). Relative protein levels normalized to the loading control tERK and to the shCtrl + SAG sample are shown above each protein band. **(C)** qPCR analysis of GLI1 and HHIP mRNA levels of Daoy cells treated as described in (B) (n = 3). Student’s t test was used for statistical analysis (**P < 0.002).

### 2.2. Genetic inhibition of CSNK1D reduces HH/GLI-activity in SMO-inhibitor resistant tumor entities driven by oncogenic GLI

To expand our investigations towards SMO-inhibitor resistant cancer cells, we turned to Ewing Sarcoma (EWS) cells, where SMO-independent GLI1 expression is driven by the EWS-FLI1 fusion oncogene [54] (Fig 2A). Further, in EWS cells GLI1 acts downstream of EWS-FLI1 as oncogene to promote proliferation and 3D spheroid growth [55]. In line with SMOi resistance of the GLI1-expressing EWS cell lines A673 and MHH-ES-1, only treatment with the GLI inhibitor HPI-1 led to reduced GLI1 protein levels, while treatment with the FDA-approved SMOi vismodegib (vismo) did not affect GLI1 expression (Fig 2B and suppl. Fig S2A)[56,57].

**Figure 2.**
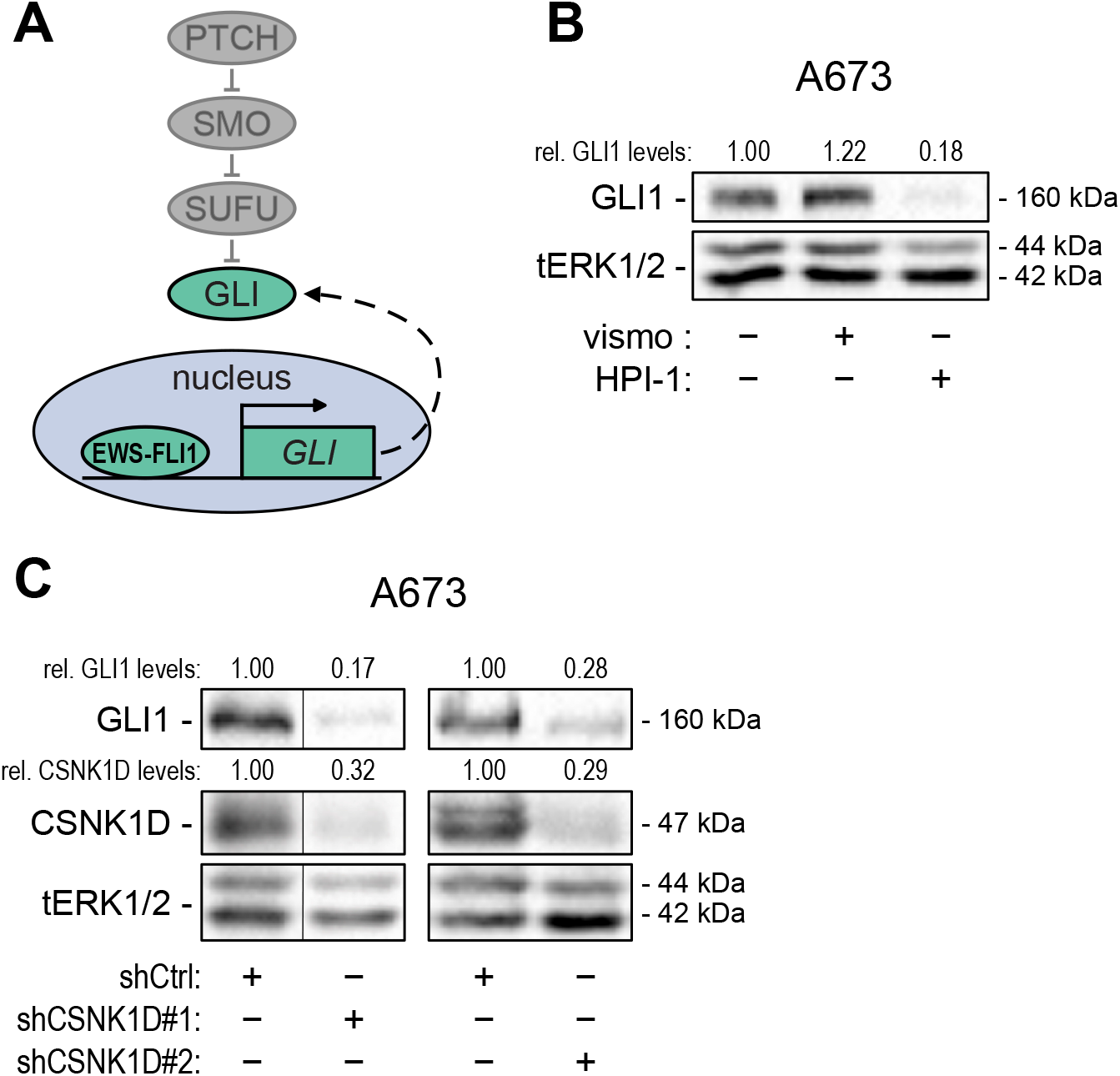
RNAi-mediated inhibition of CSNK1D interferes with oncogenic HH/GLI signaling in SMOi resistant Ewing Sarcoma cells. **(A)** Simplified schematic illustration of non-canonical, SMO-independent HH/GLI signaling in Ewing Sarcoma with EWS-FLI1-driven transcription of *GLI1*. **(B)** Representative Western blot analysis of GLI1 in A673 cells treated with vismodegib [1 μM] or HPI-1 [20 μM]. **(C)** Representative Western blot analysis of GLI1 in A673 cells lentivirally transduced with shCSNK1D (#1, #2) or control shRNA (shCtrl). Relative quantification of Western blot bands was conducted via densitometric image analysis using Image Lab 5.0 software (Bio-Rad, Vienna, Austria). Relative protein levels normalized to the loading control tERK and to the Ctrl sample are shown above each protein band.

To investigate whether CSNK1D regulates GLI activity also in SMO-independent cells, we performed shRNA-mediated knockdown of CSNK1D in A673 cells. As shown in Fig. 2C and suppl. Fig. S2B, genetic perturbation of CSNK1D reduced GLI1 protein levels, suggesting that CSNK1D also regulates SMO-independent GLI activity. To support these results, we inactivated CSNK1D by CRISPR-Cas9 in the EWS cell line MHH-ES-1 and confirm that deletion of CSNK1D results in reduced GLI1 protein levels (suppl. Fig S2C).

### 2.3. Pharmacological targeting of CSNK1D inhibits HH/GLI signaling in both canonical and non-canonical settings

The critical role of CSNK1D in SMO-dependent and SMO-independent GLI activation prompted us to analyze the possible therapeutic effect of CSNK1D inhibitors by targeting oncogenic GLI transcription factors. In a collaborative approach, we performed a compound screen cascade in A549 non-small cell lung carcinoma (NSCLC) cells under proliferating and non-proliferating conditions to select for compounds interfering with quiescent and tumor initiating cancer cell properties. This approach identified a novel, highly effective CSNK1D inhibitor, termed CK1D008 (suppl. Fig. S3A). To investigate whether chemical perturbation of CSNK1D can interfere with HH/GLI signaling, we treated HH-responsive Daoy cells with CK1D008 and for comparison with the known CSNK1D inhibitor SR-3029 [58]. As shown in Fig. 3A, both CSNK1D inhibitors potently inhibited the activation of GLI1 mRNA expression in SAG-treated Daoy cells with IC50 values in the nanomolar range (Fig 3A). Likewise, both inhibitors decreased SAG-induced GLI1 protein levels in Daoy cells in a dose-dependent manner (Fig 3B). Neither the protein levels of CSNK1D nor those of GLI2 were affected by pharmacological inhibition of CSNK1D (suppl. Fig S3B). Consistent with the proliferative role of HH/GLI in Daoy cells, both CSNK1D inhibitors also reduced cell proliferation and viability in a dose-dependent manner (suppl. Fig S3C).

**Figure 3.**
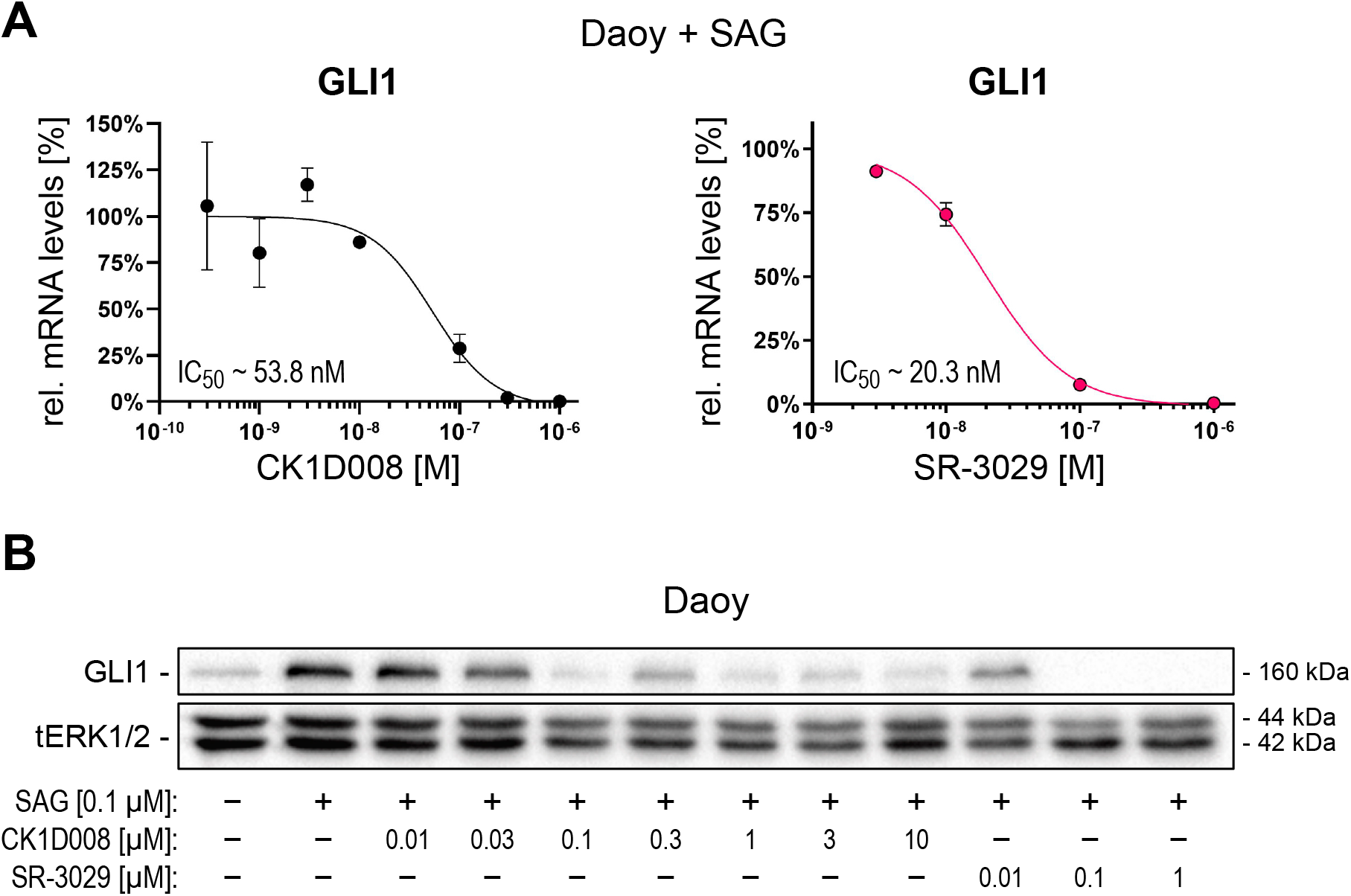
Pharmacological targeting of CSNK1D inhibits SMO-dependent GLI activation. **(A)** mRNA expression levels of GLI1 were analyzed by qPCR and expressed as percentage relative to the control sample (n = 3). **(B)** Representative Western blot analysis of GLI1 protein levels in Daoy cells treated with SAG [100 nM] and increasing concentrations of CK1D008 [0.01 – 10 μM] or SR-3029 [0.01 – 1 μM].

Furthermore, to mimic SMO inhibitor resistance, we generated Daoy cells with an shRNA-mediated knockdown of the GLI inhibitor SUFU [59], resulting in SMO-independent activation of GLI1 expression (Fig 4A, B). Of note, while vismodegib treatment failed to reduce GLI1 expression in Daoy-shSUFU cells, chemical targeting of CSNK1D with CK1D008 or SR-3029 efficiently reduced GLI1 protein expression and HH target gene expression in this SMO-independent model of GLI activation (Fig 4B and suppl. Fig S4A). To corroborate that CSNK1D inhibition is a potent strategy to block GLI activation in SMOi-resistant settings, we performed chemical inhibition of CSNK1D in A673 and MHH-ES-1 Ewing sarcoma cells both showing SMO-independent GLI1 expression. As shown in Fig. 4C and suppl. Figure 4B, pharmacologic inhibition of CSNK1D with CK1D008 or SR-3029 efficiently reduced GLI1 expression in both cell lines. By contrast, vismodegib treatment did not affect GLI1 expression [60].

**Figure 4.**
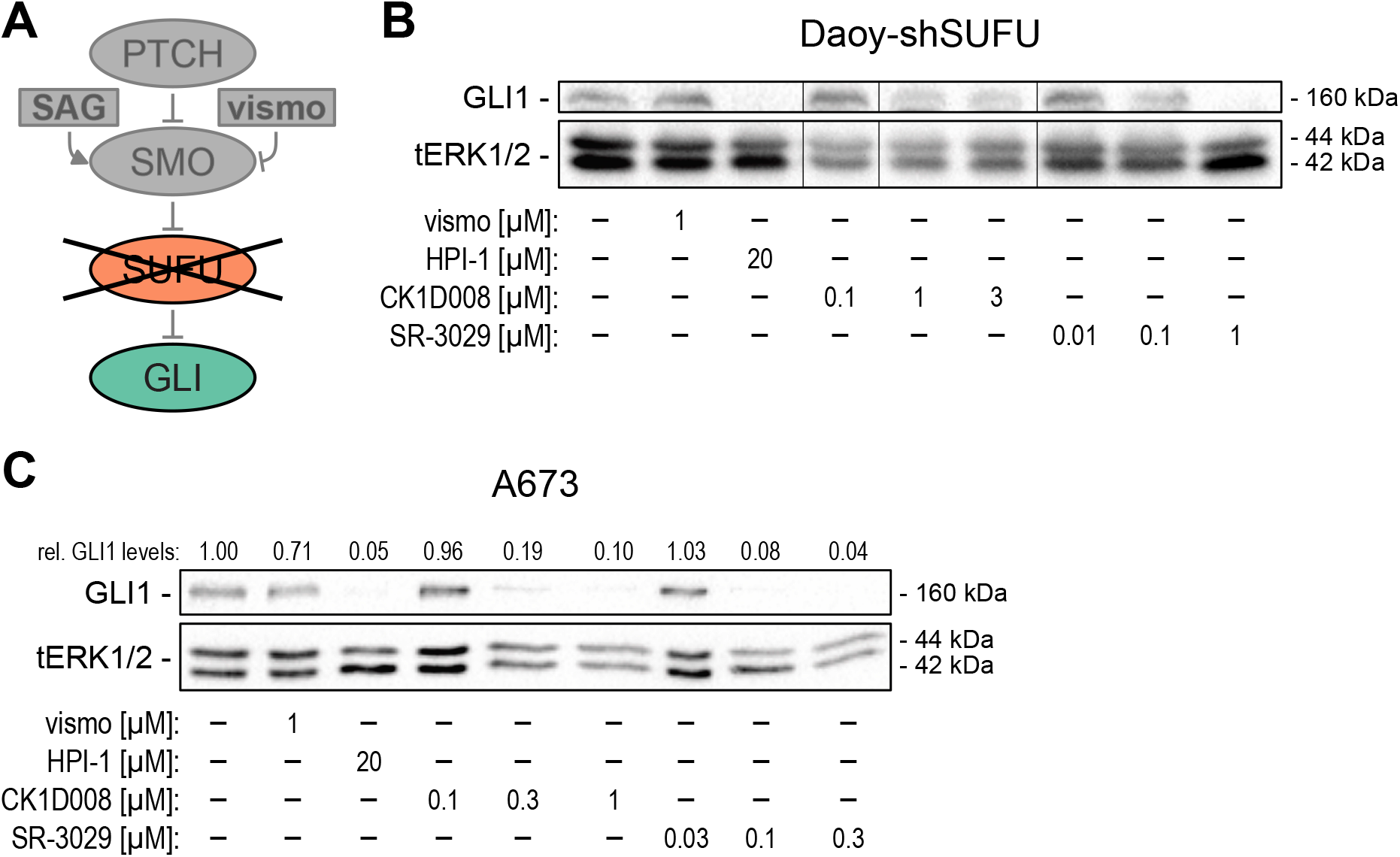
Pharmacological targeting of CSNK1D inhibits oncogenic HH/GLI signaling in SMOi resistant cells. **(A)** Simplified schematic illustration of SMO independent activation of HH/GLI signaling in Daoy medulloblastoma cells caused by the loss of the negative HH/GLI pathway regulator SUFU. **(B)** Representative Western blot analysis of GLI1 protein levels in Daoy-shSUFU cells treated with vismo [1 μM], HPI-1 [20 μM], CK1D008 [0.1 – 3 μM] or SR-3029 [0.01 – 1 μM]. **(C)** Representative Western blot analysis of GLI1 in A673 cells treated with vismo [1 μM], HPI-1 [20 μM], CK1D008 [0.1 – 1 μM] or SR-3029 [0.03 – 0.3 μM]. Relative quantification of Western blot bands was conducted via densitometric image analysis using Image Lab 5.0 software (Bio-Rad, Vienna, Austria). Relative protein levels normalized to the loading control tERK and to the Ctrl sample are shown above each protein band.

### 2.5. Targeting the CSNK1D-GLI axis inhibits CSC-like characteristics in vitro and in vivo

Clonogenic and self-renewing spheroid growth in 3-dimensional *in vitro* cultures and the initiation of *in vivo* tumor growth are considered characteristics of tumor-initiating cells [60]. To functionally addressed a putative role of the CSNK1D-GLI axis in TICs, we first performed genetic and pharmacologic targeting of CSNK1D and measured its impact on the 3D spheroid growth properties of GLI1-expressing A673 cells. Of note, shRNA-mediated knockdown of CSNK1D significantly diminished the clonogenic anchorage-independent growth capacity of A673 cells in 3D cultures (Fig 5A), while it did not affect anchorage-dependent growth under 2-D culture conditions (suppl. Fig S5A). Accordingly, pharmacological inhibition of CSNK1D with CK1D008 or SR-3029 both drastically reduced clonogenic spheroid growth in 3-D cultures at concentrations that had no impact on cell growth under planar 2-D conditions (Fig 5B and suppl. Fig S5B). In addition, the inhibitory effect of CK1D008 on clonal growth was also evaluated in HH/GLI-dependent pancreatic (PANC1), NSCLC (A549), glioma (LNT-229) and CRC (HT29, HCT15, HCT116) tumor cells [6,60–62]. To this end, tumor cells were pre-treated with inhibitor for 48h and viable cells were seeded at limiting dilutions in an anchorage-dependent colony formation assay. CK1D008 efficiently reduced the colony formation ability of all tested cell lines at concentrations as low as 0.1-0.3 μM (suppl. Fig S5C).

**Figure 5.**
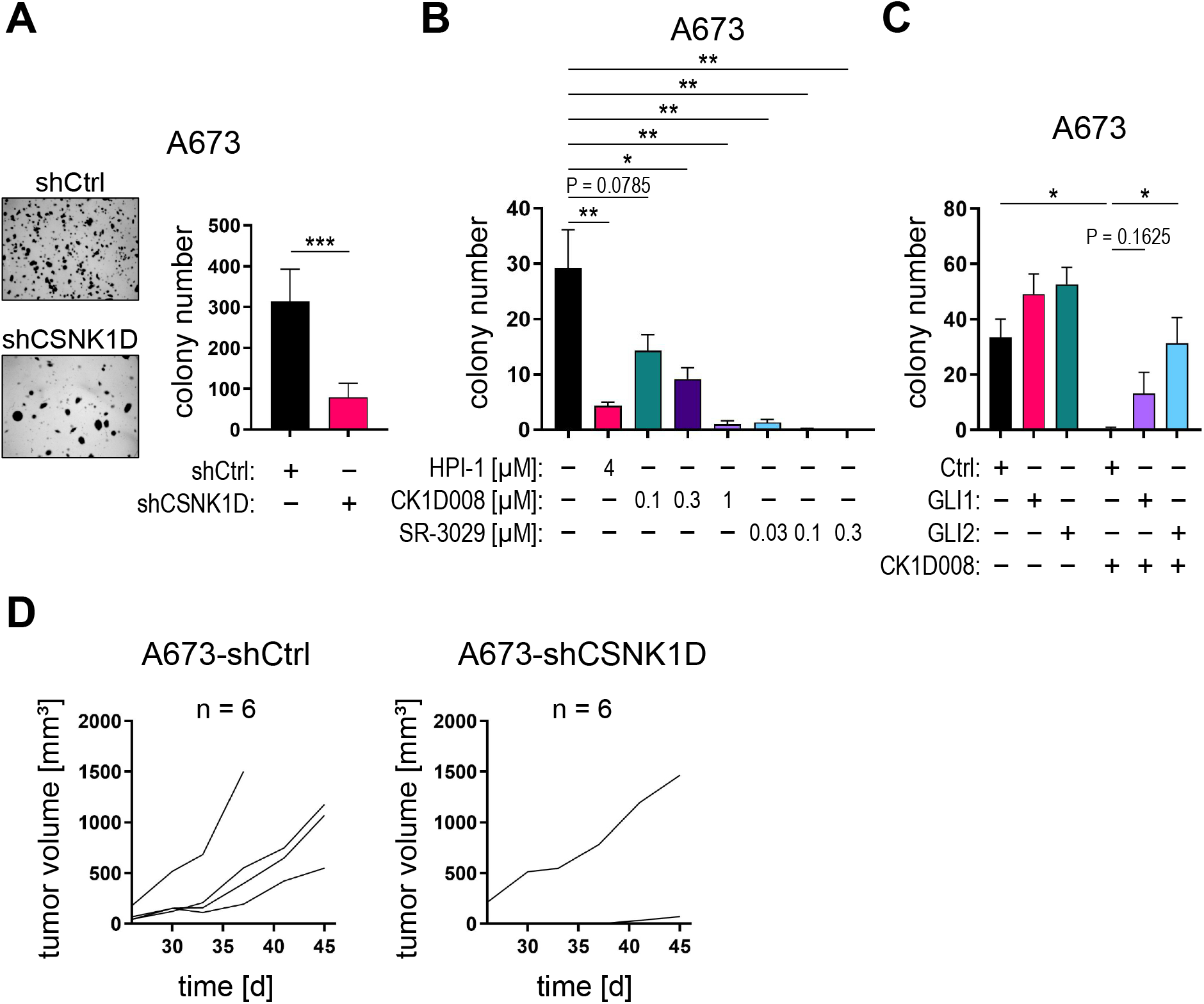
Targeting CSNK1D reduces GLI-dependent 3D tumor spheroid formation *in vitro* as well as tumor engraftment *in vivo*. **(A)** A673 cells were lentivirally transduced with shCSNK1D or control shRNA (shCtrl) and cultured under anchorage-independent conditions. Representative images of formed 3D spheroid colonies (left panel). The number of 3D spheroid colonies was counted (right panel) (n = 8). **(B)** A673 cells were cultured under anchorage-independent conditions and treated with the GLI inhibitor HPI-1 [4 μM], CK1D008 [0.1 – 1 μM] or SR-3029 [0.03 – 0.3 μM]. The data represent the mean of six 3D-culture experiments. **(C)** A673 cells were retrovirally transduced with GLI1, active GLI2 (GLI2act) or control overexpression constructs, cultured under anchorage-independent conditions and treated with CK1D008 [0.3 μM]. The number of 3D spheroid colonies was counted (n = 4 experiments). **(D)** A673 cells were lentivirally transduced with shCSNK1D or control shRNA (shCtrl) and engrafted in the flanks of NSG mice (n = 6 mice each). Tumor volume was measured every 3-4 days. Student’s t test was used for statistical analysis (*P < 0.035; **P < 0.002; ***P < 0.0002).

GLI1 represents an important driver gene of proliferation and spheroid growth in Ewing Sarcoma cells [29]. To further support that inhibition of CSNK1D decreases TIC properties via GLI1 inhibition, we performed a CRISPR/Cas-mediated GLI1 knockout in A673 cells (suppl. Fig S5D) and analyzed GLI1-deficient A673 cells for clonogenic spheroid growth in 3-D and planar growth in 2-D cultures. In line with our results on CSKN1D targeting, genetic deletion of GLI1 selectively abolished clonogenic spheroid formation in 3-D, while it did not affect the 2-D growth properties (suppl. Fig S5E, F), suggesting that CSNK1D encodes a crucial positive regulator of oncogenic GLI1 in TIC.

To evaluate whether enforced overexpression of the GLI activator proteins GLI1 and GLI2 can attenuate the anti-tumor effect caused by CSNK1D inhibition, we overexpressed in A673 cells HA-tagged GLI1 and an active form of GLI2 [63], respectively (suppl. Fig S5G) and treated GLI1, GLI2 or empty-vector control cells with CK1D008. In line with GLI acting downstream of CSNK1D, GLI1 and GLI2-overexpression conferred at least partial resistance to CSNK1D inhibitor treatment compared to control cells (Fig 5C).

To investigate whether CSNK1D is required for GLI-driven tumor initiation *in vivo,* we performed xenograft experiments using GLI1-dependent A673 and GLI1-dependent PANC1 pancreatic cancer cells [60] with concomitant inhibition of CSNK1D. As shown in Fig. 5D, CSNK1D knockdown in A673 cells severely impaired their engraftment ability compared to control cells transduced with non-targeting shRNA. Similarly, pretreatment of GLI1-dependent PANC1 cells with a low concentration of CK1D008 [0.3 μM] effectively abrogated the engraftment capacity of PANC1 cells (suppl. Fig S5H). These data strongly support a model where CSNK1D regulates the oncogenic activity of activator GLIs in tumor-initiating cells independent of SMO function.

## 3. Discussion

Hedgehog/GLI signaling has been associated with many human malignancies and aberrant pathway activation has been discovered in cancer stem cells with a critical role in tumor initiation, malignant growth, metastasis, and relapse [3,4,7,8,64]. Pharmacological targeting of oncogenic HH/GLI is therefore considered a promising therapeutic strategy. Three SMO inhibitors, vismodegib, sonidegib and glasdegib, have been approved for the treatment of locally advanced and metastatic basal cell carcinoma and for acute myeloid leukemia, respectively [20,21,23,57]. Even though these SMO inhibitors demonstrate remarkable therapeutic efficacy, treatment can cause frequent and severe side effects and result in acquired drug resistance. Additionally, several tumor entities are driven by SMO-independent GLI activity and thus display *a priori* SMOi resistance [25,26,30,54]. Since, the GLI transcription factors can integrate with other oncogenic signaling cascades and promote cancer progression and malignant properties of cancer (stem) cells [5,6,34,38–40,60,62,65], oncogenic GLI transcription factors represent attractive therapeutic targets for patients with acquired and *a priori* resistance to SMOi. Although some previous studies have shown that direct targeting of oncogenic GLI proteins is feasible and promising [66,67], small molecule-mediated direct inhibition of transcription factors is generally considered to be very challenging. The identification of druggable key regulators of GLI activity such as readily targetable kinases promoting the oncogenic activity of GLI transcription factors is therefore a critical requirement and of high medical relevance and need.

In this study, we identify CSNK1D as a positive and druggable regulator of oncogenic GLI activity, both in canonical as well as SMO-independent settings of GLI activation. Furthermore, we describe the identification of the novel CSNK1D inhibitor CK1D008 and show that pharmacological targeting of CSNK1D with potent small molecule inhibitors including CK1D008 and SR-3029 [68] abolishes oncogenic HH/GLI signaling in distinct cancer entities with either SMO-dependent or SMO-independent oncogenic GLI activity. Of note, we provide evidence that the CSNK1D-GLI axis selectively promotes hallmarks of TICs including clonogenic growth in 2D- and 3D cultures and engraftment in immunocompromised mouse models, while not affecting the proliferation of non-TICs under standard 2D culture conditions. Therapeutic targeting of CSNK1D may therefore represent a promising approach to eradicate highly malignant GLI-driven TICs. Considering recent data on the immunosuppressive role of TICs/CSCs in several cancer entities [69] as well as the regulation of immunosuppressive factors by GLI [70–73] combination therapy with CSNK1D and immune checkpoint inhibitors is an attractive therapeutic strategy to be evaluated in follow-up pre-clinical studies.

Casein kinase 1 family members interact with several oncogenic pathways such as the Hedgehog, Hippo and Wnt/beta catenin pathway, and their aberrant regulation has been closely linked to several human malignancies [74]. Small molecule inhibitors of casein kinase 1 family members have been developed, such as the CSNK1D/E inhibitor SR-3029 which is a potent ATP-competitor with high specificity for CSNK1D and CSNK1E [68]. SR-3029 displays striking therapeutic efficacy in triple negative as well as HER2^+^ breast cancer models with Wnt/β-catenin involvement [58]. Given the documented oncogenic role of HH/GLI in breast cancer development [75–77], it is possible that the therapeutic activity of this CSNK1 inhibitor also relies at least in part on the GLI inhibitory activity described in our study.

Mechanistically, we propose that CSNK1D positively regulates the stability of oncogenic GLI proteins. In this context it is noteworthy that Shi *et al.* (2014) have shown that members of the CSNK1 family phosphorylate GLI transcription factors at PKA-independent sites, thereby disrupting the interaction of GLI with SPOP. SPOP negatively regulates GLI activity by enhancing the proteasomal degradation of GLI proteins [50]. In line with these data, we observed that chemical inhibition of the proteasome machinery at least partially reversed the repressive effect of CSNK1D drugs on GLI protein expression levels (data not shown).

To address the oncogenic role of the CSNK1D-GLI axis in stem-like TICs, we performed 3D growth assays to monitor clonal spheroid growth, which is considered a hallmark of tumorigenic TICs [60]. Genetic inhibition of CSNK1D was sufficient to abrogate colony formation in Ewing sarcoma cells. Pharmacological targeting of CSNK1D by CK1D008 and SR-3029 selectively decreased colony formation at concentrations sufficient to abrogate HH pathway activity, while not affecting cell proliferation in planar 2D culture settings. This suggests that CSNK1D-GLI is preferentially required for the expansion of TICs rather than non-CSCs. In agreement with this notion, we show that genetic inhibition of CSNK1D diminishes the engraftment and tumor initiation capacity of GLI-dependent A673 Ewing sarcoma cells in immunodeficient mice. The tumor-initiating role of CSNK1D is further supported by our findings that pharmacological inhibition of CSNK1D abrogates the tumor initiation and *in vivo* engraftment capacity of GLI-dependent PANC1 cells [60].

In summary, we identified CSNK1D as a novel positive regulator of oncogenic GLI transcription factors in tumor-initiating cancer cells. This study provides a basis for the development and use of selective CSNK1D inhibitors to abrogate HH/GLI signaling in SMOi-sensitive and SMOi-resistant settings, which is an important step towards the development of novel oncology drugs targeting GLI transcription factors to eliminate highly malignant cancer stem cells.

## 4. Materials and Methods

### Cell lines and reagents

Daoy medulloblastoma cells (ATCC HTB-186), Ewing Sarcoma cell lines A673 (ATCC CRL-1598) and MHH-ES-1 (DSMZ ACC 167) were used for chemical and genetic manipulation of CSNK1 and HH signaling pathway components. The following chemicals were used: Smoothened agonist SAG (Selleckchem, Houston, TX, United States), GDC-0449 (vismodegib; LC Laboratories, Woburn, MA, United States), HPI-1 (Sigma-Aldrich, St Louis, MO, United States), SR-3029 (Axon Medchem, Groningen, Netherlands) and CK1D008 (4SC AG, Planegg-Martinsried, Germany). For the analysis of HH/GLI activity, Daoy cells were kept confluent for at least 48h and starved in 0.5% FBS (Sigma-Aldrich) overnight prior to stimulation with 100 nM SAG. Chemicals or control solvents were added 2h prior to SAG stimulation. Daoy cells were cultured in MEM (Sigma-Aldrich) supplemented with 10% FBS (Sigma-Aldrich) and antibiotics (Penicillin-Streptomycin, Sigma-Aldrich). A673 cells were cultured in DMEM (Sigma-Aldrich) and MHH-ES-1 cells in RPMI-1640 (Sigma-Aldrich) supplemented with 10% FBS and antibiotics and treated at confluency as indicated in the text. HEK293FT cells were cultured in DMEM supplemented with 10% FBS, antibiotics, L-Glutamin (Sigma-Aldrich) and non-essential amino acids solution (Thermo Fisher Scientific, Waltham, MA, United States) and used for transfection experiments when they reached ~80% confluency.

### Screening cascade

A549 and PANC1 cells were seeded under proliferating (10%) FBS and non-proliferating (0.2%) FBS conditions and treated with screening compounds for 48 h and analyzed with crystal violet staining. Compounds potently reducing the cell numbers under 0.2% FBS and a factor of 10 under proliferating conditions were selected for further analysis.

### Cell proliferation and anchorage-dependent and -independent growth assays

Cell proliferation and viability were determined by the AlamarBlue™ assay. Cells were seeded in 96-well plates at an appropriate density depending on their growth properties to ensure that confluency would not be reached during the experiment. Cell viability was determined at indicated time points by adding 1/10 volume AlamarBlue solution (Biorad, Hercules, CA, United States) to each well, followed by measuring fluorescence with an excitation of 560 nm and an emission of 590 nm, or absorbance at 570 nm and 600 nm. The percentage of viable cells was normalized to the number of viable cells in the respective control.

For anchorage-independent three-dimensional (3D) spheroid colony growth cultures, 1×10^4^ cells were seeded in a 12-well plate as described in [65].

For anchorage-dependent limited dilution colony formation assay, tumor cells were pre-treated with compounds for 48 h at 0.2% FBS. 200 viable cells per well were seeded into 6-well plates and incubated without compound for 9-11 days followed by crystal violet staining and colony counting. To evaluate the potential irreversible toxic effect of compound pre-treatment, cells were incubated without compound for another 24h and analyzed for viability. Only compounds that were not toxic in this assay were considered as colony formation inhibitors.

### In vivo experiments

For *in vivo* experiments, NOD-SCID IL2Rgamma^-/-^ (NSG) mice were kept in individually ventilated cages (IVC) under specific pathogen-free conditions (SPF). All animal experiments were performed in compliance with the national requirements and regulations. The A673 allograft assay was performed by injecting 1×10^6^ A673 cells per 100μL 25% Matrigel (BD Biosciences, Franklin Lakes, NJ, United States) subcutaneously into the flanks of NSG mice. Tumor volume was measured every 3-4 days with a caliper and calculated according to the formula V=4/3*π*length/2*width/2*height/2. For PANC1 engraftment experiments, PANC1 cells were treated for 48 h prior to engraftment with DMSO or 0.3 μM CK1D008 and analyzed for viability. 1×10^6^ viable PANC1 cells per mouse were inoculated into BalbC Nu/Nu athymic mice and monitored for tumor growth. Tumor growth was calculated according to the formula V= (a*b*b)/2, where a and b correspond to the longest and shortest diameter of the engraftment.

### RNA isolation and quantitative PCR (qPCR)

Total RNA was isolated using TRI reagent (Sigma-Aldrich) according to the manufacturer’s protocol followed by LiCl precipitation. cDNA was synthesized with the M-MLV Reverse Transcriptase (Promega, Madsion, WI, United States) and qPCR was done on the Rotor-Gene Q instrument (Qiagen, Hilden, Germany) using the GoTaq qPCR Mastermix (Promega). The qPCR primers used are listed in Supplementary Table S1.

### Western blot analysis

After genomic modification or small molecule inhibitor treatment, cell samples were harvested and lyzed in Laemmli buffer [78] supplemented with phosphatase and protease inhibitors. Proteins were separated by SDS-PAGE and then blotted onto hybond ECL membranes. Antibodies used are listed in the Supplementary Table S2.

### RNA interference and overexpression constructs

For RNA interference, two CSNK1D targeting shRNA constructs were selected from the Mission TRC shRNA library (TRCN0000023769, named shCSNK1D#1; TRCN0000361946, named shCSNK1D#2; Sigma-Aldrich); non-targeting scrambled shRNA served as control (shc002, named shCtrl, Sigma-Aldrich). SMO-inhibitor resistant Daoy cells with shRNA-mediated knockdown of SUFU had been generated in our lab using the TRCN0000019466 construct (Sigma-Aldrich) [52].

For overexpression of GLI transcription factors, GLI1 and active GLI2 construct (GLI2act) [63] were HA-tagged and cloned into an empty pMSCV-puro vector using the Gibson Assembly method. Retroviral transduction experiments were performed as described in [79]. Transduced cells were selected for puromycin resistance prior to further analysis.

### CRISPR-mediated knockout

Lentiviral single guide (sg) RNA expression vector targeting CSNK1D (#GSGH11838-246527777) and the corresponding non-targeting control (#GSG11811) were purchased from Dharmacon (Horizon Discovery, Waterbeach, United Kingdom). For expression of the Cas9 protein the lentiCas9-Blast expression vector was used (Addgene plasmid #59262; http://n2t.net/addgene:52962; RRID:Addgene_52962) [80]. Cells were first transduced with the lentiCas9-Blast vector, selected for blasticidin resistance and afterwards lentivirally transduced with the CSNK1D-targeting sgRNA construct conferring puromycin resistance. Knockout efficacy was determined by target-locus sequencing of genomic DNA isolated using the DNA Blood Mini Kit (Qiagen, Hilden, Germany). Knockout efficacy was calculated using TIDE (https://tide.deskgen.com) [81].

A GLI1 sgRNA targeting sequence was designed using GPP sgRNA designer (https://portals.broadinstitute.org/gpp/public/analysis-tools/sgrna-design), a sgRNA targeting eGFP was used as control. sgRNA sequences are listed in Supplementary Table S3. sgRNA oligos were cloned into the lentiCRISPRv2 vector (Addgene plasmid #52961; http://n2t.net/addgene:52961; RRID:Addgene_52961) [80]. Cells were transduced with either the GLI1-targeting or the control construct and selected for puromycin resistance as described in [79]. GLI1 knockout efficacy was calculated using TIDE (https://tide.deskgen.com) [81].

To isolate GLI1 knockout cells, sgGLI1-transduced cells were seeded at low density in a 96-well plate and cultured in A673-conditioned media cleared by filtration through 0.45 μm filters. Clonal colonies were expanded for further experiments.

### Statistical analysis

Statistical analysis and graph design were done using GraphPad Prism 8 software (GraphPad Software, San Diego, CA, USA).

## 5. Conclusions

In this study, we identified CSNK1D as critical positive regulator of oncogenic GLI activity and demonstrate that targeting CSNK1D interferes with the malignant properties of GLI-dependent tumor-initiating cells. Of note, genetic as well as pharmacologic inhibition of CSNK1D efficiently represses oncogenic GLI activity in cancer cells resistant to FDA-approved HH pathway inhibitors targeting the essential HH effector SMO. Targeting CSNK1D with compounds such as SR-3029 and CK1D008 may therefore be a promising future strategy to treat cancer patients with acquired or *a priori* resistance to SMO-inhibitors and to hopefully reduce severe side effects known to be caused by established anti-SMO drugs.

## Author Contributions

Conceptualization, E.P., W.G., S.H. and F.A.; data validation, formal analysis, investigation, E.P., W.G., T.P. S.A.; methods and resources, A.R., S.G., F.R.; medicinal chemistry, conceptualization, compound screening and biological evaluation of CK1D008, M.Z., S.M. and S.H.; data curation, E.P., T.P., S.A. and F.V.; manuscript writing E.P., D.E., S.G., S.H. and F.A.; visualization, E.P., D.E. and S.A.; supervision, project administration and funding acquisition F.A.; All authors have read and agreed to the published version of the manuscript.

## Acknowledgments

The authors thank Sabine Siller and Philip Markus Wolf for their excellent technical support.

## Abbreviations

BCC: Basal cell carcinoma
CSNK1: Casein kinase 1
GLI: Glioma-associated oncogene homolog
HH: Hedgehog
HHIP: Hedgehog interacting protein
PTCH: Patched
SAG: Smoothened Agonist
SMO: Smoothened
SPOP: Speckle-type POZ protein
SUFU: Suppressor of Fused

