## Supplementary figures and images for "Casein kinase 1D encodes a novel drug target in Hedgehog-GLI driven cancers and tumor-initiating cells resistant to SMO inhibition"

### suppl Figures

Peer et al. Supplementary Figures

**Figure S1**

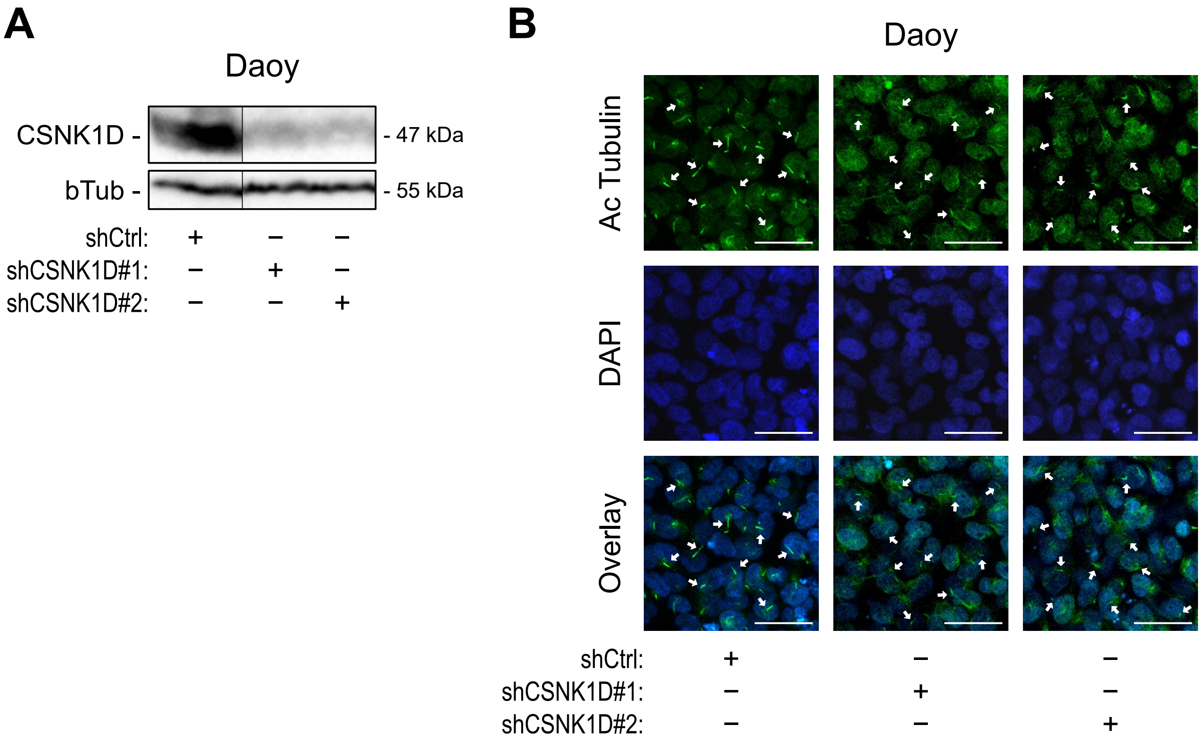

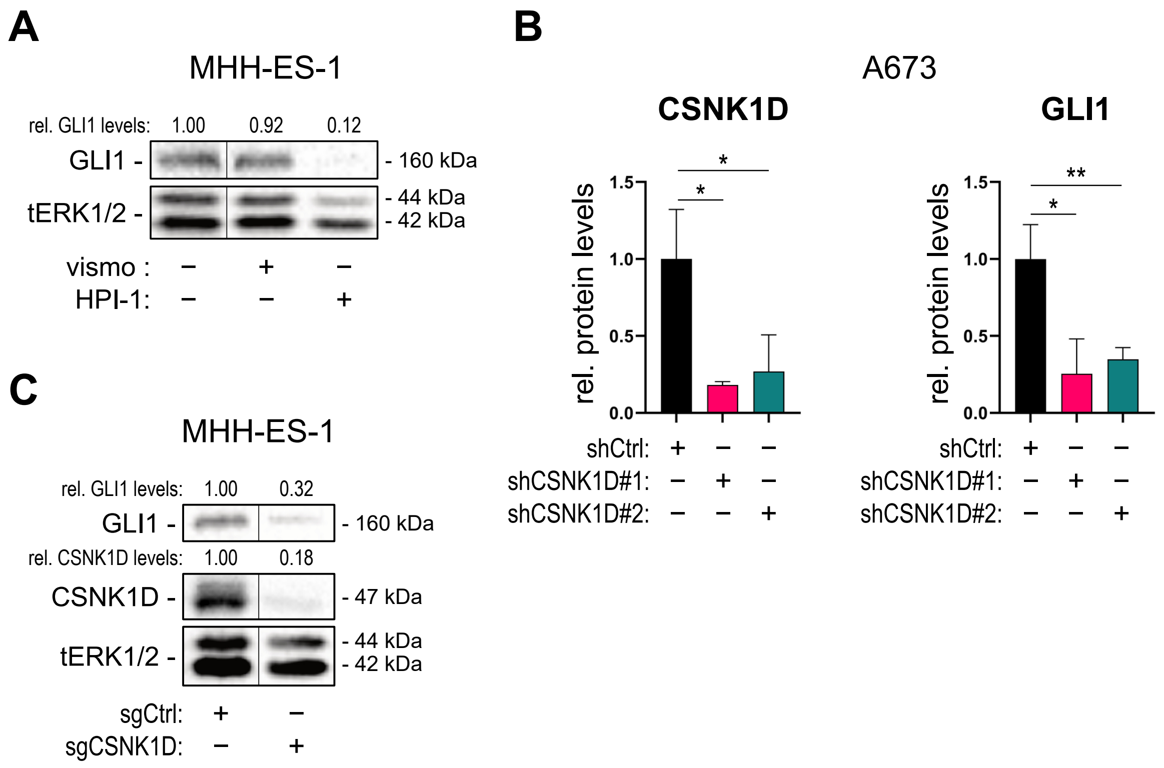

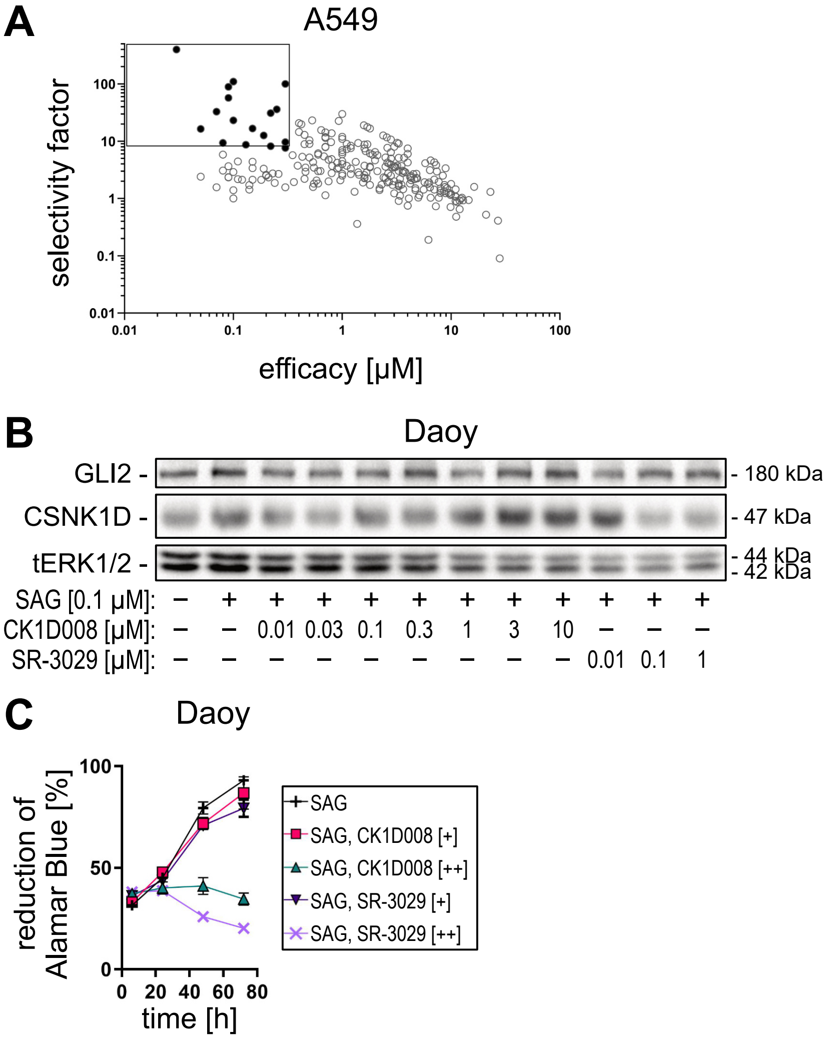

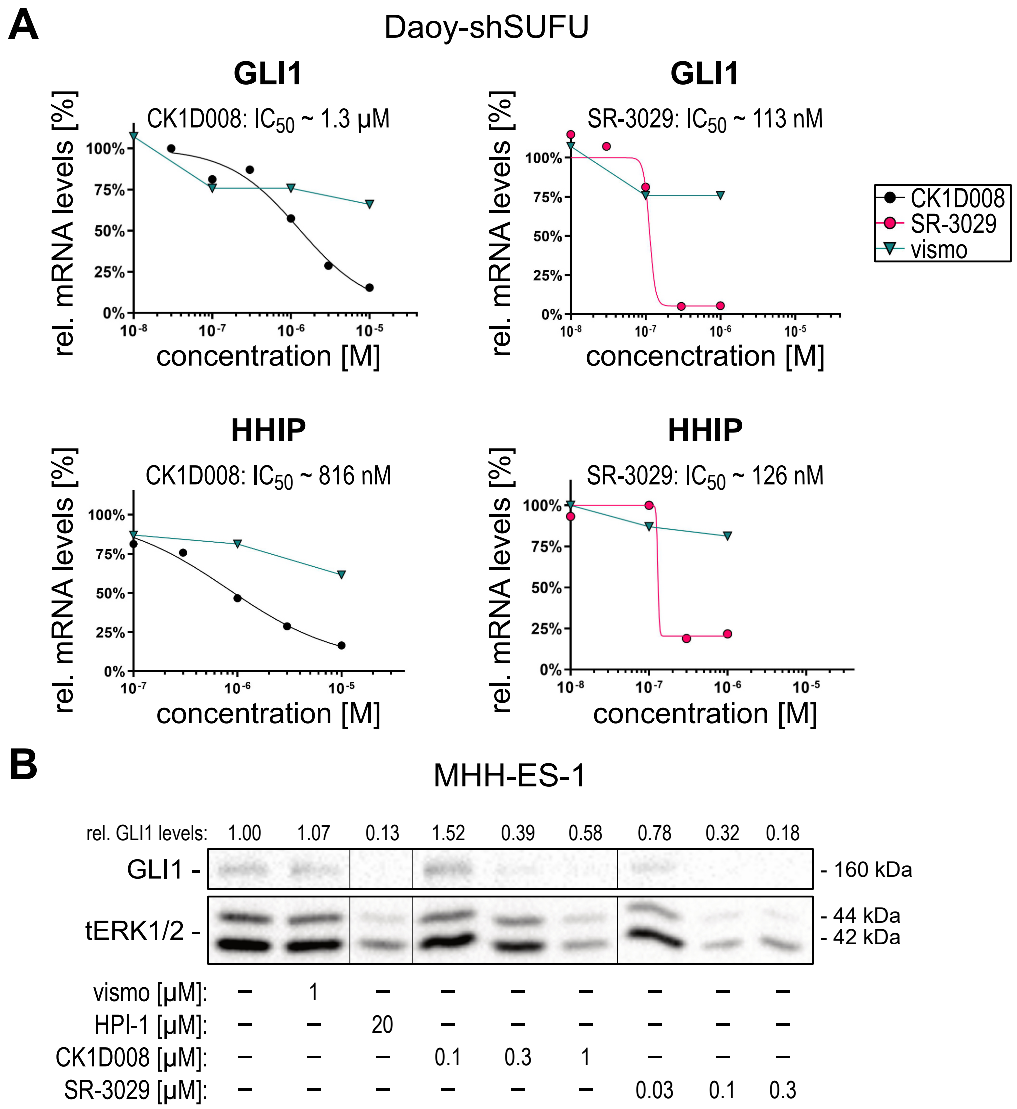

Peer et al. Supplementary Figures

#### Figure S5

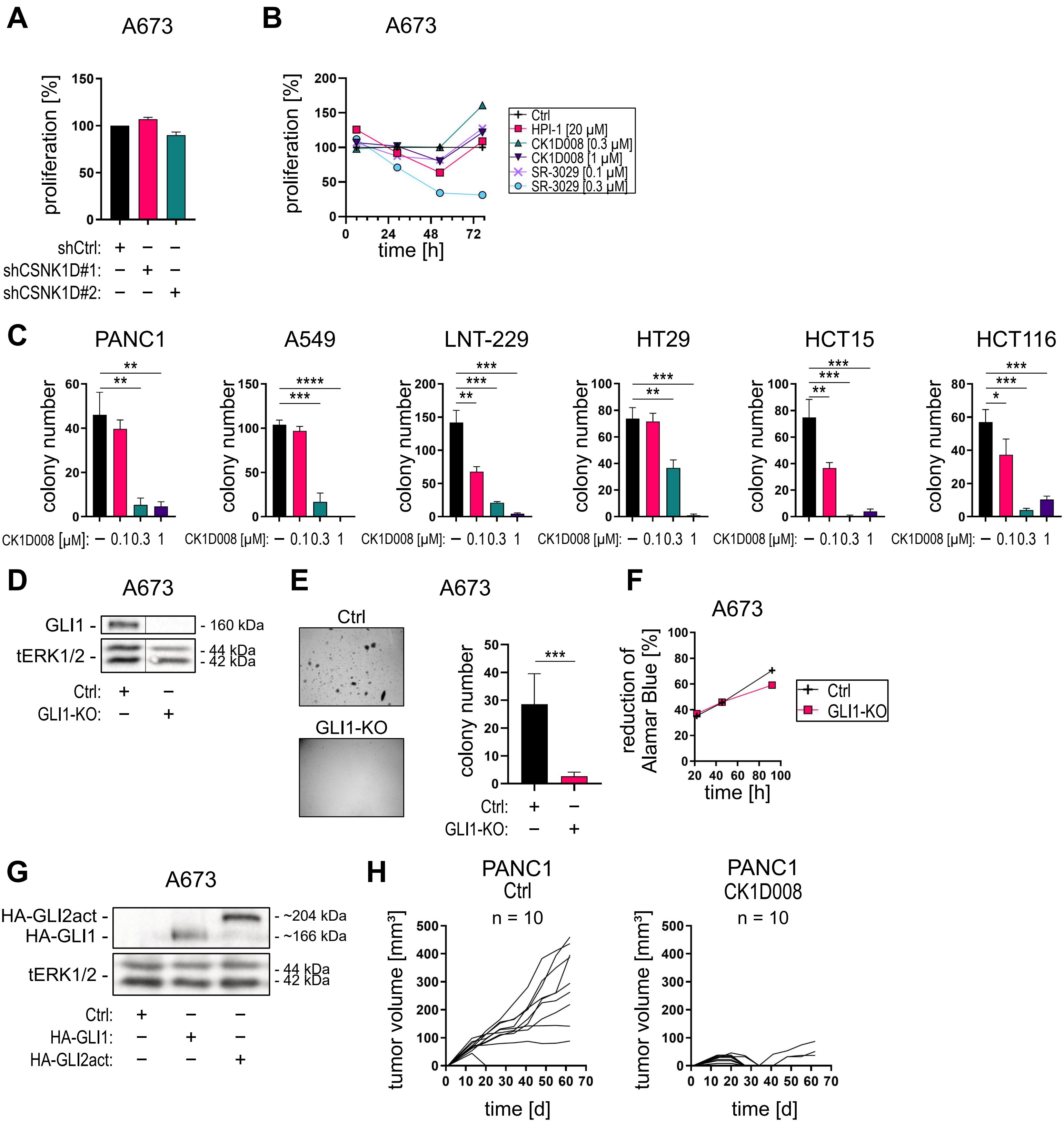
