## Supplementary material for "Casein kinase 1D encodes a novel drug target in Hedgehog-GLI driven cancers and tumor-initiating cells resistant to SMO inhibition": suppl figure legends

Peer et al. – SUPPLEMENTARY FIGURES – legends:

**Suppl. Fig. S1 Targeting CSNK1D does not interfere with ciliogenesis in Daoy cells. (A)** Representative Western blot analysis of CSNK1D in Daoy cells lentivirally transduced with shCSNK1D (#1, #2) or control shRNA (shCtrl). **(B)** Confocal imaging of primary cilia in Daoy cells lentivirally transduced with shCSNK1D (#1, #2) or control shRNA (shCtrl). For visualization of primary cilia (indicated by white arrows) Daoy cells were stained with antibodies against acetylated tubulin as described previously [1]. Scale bars: 50 µm.

**Suppl. Fig. S2 Targeting CSNK1D interferes with oncogenic HH/GLI signaling in SMOi resistant Ewing Sarcoma cells. (A)** Representative Western blot analysis of GLI1 in MHH-ES-1 cells treated with vismodegib [1 µM] or HPI-1 [20 µM]. **(B)** Quantification of relative GLI1 protein levels in A673 cells lentivirally transduced with shCSNK1D (#1, #2) or control shRNA (shCtrl) (n = 3). **(C)** Representative Western blot analysis of GLI1 in MHH-ES-1 cells lentivirally transduced with sgCSNK1D or sgCtrl. Quantification of Western blot bands was conducted via densitometric image analysis using Image Lab 5.0 software (Bio-Rad). Relative protein levels normalized to the loading control total ERK (tERK) and to the Ctrl condition are indicated above each protein band. Student's t test was used for statistical analysis (*P < 0.035; **P < 0.002).

**Suppl. Fig. S3 Effects of pharmacological targeting of CSNK1D on the proliferation of A549 and Daoy cells (A)** Cellular compound screening cascade in A549 cells. Compounds (black dots) with an $IC_{50}$ below 0.3 µM under non-proliferating conditions (= efficacy) and a selectivity factor of at least 10, such as CK1D008, were selected for further evaluation. **(B)** Representative Western blot analysis of GLI2 and CSNK1D protein levels in Daoy cells treated with SAG [100 nM] and increasing concentrations of CK1D008 [0.01 – 10 µM] or SR-3029 [0.01 – 1 µM]. **(C)** Daoy cells were treated with SAG [100 nM] and increasing concentrations of CK1D008 ([0.3 µM] (+), [3 µM] (++)] or SR-3029 ([0.1 µM] (+), [1 µM] (++)) and proliferation was assessed in AlamarBlue assays.

**Suppl. Fig. S4 Targeting of CSNK1D reduces HH target gene expression in SMOi resistant cell lines. (A)** CK1D008 and SR-3029 reduce HH/GLI target gene expression in Daoy-shSUFU medulloblastoma cells with $IC_{50}$ values in the low micromolar or nanomolar range, respectively. mRNA expression levels of GLI1 and HHIP were analyzed by qPCR and expressed as percentage relative to the control condition. **(B)** Representative Western blot analysis of GLI1 in MHH-ES-1 cells treated with vismo [1 µM], HPI-1 [20 µM], CK1D008 [0.1 – 1 µM] or SR-3029 [0.03 – 0.3 µM]. Relative quantification of Western blot bands was conducted via densitometric image analysis using Image Lab 5.0 software (Bio-Rad). Relative protein levels normalized to the loading control total ERK (tERK) and to the Ctrl condition are indicated above each protein band.

**Suppl. Fig. S5 Pharmacologic and genetic inhibition of CSNK1D selectively abrogates clonal growth and impairs *in vivo* engraftment of GLI-dependent PANC-1 cancer cells. (A)** A673 cells were lentivirally transduced with shCSNK1D (#1, #2) or control shRNA (shCtrl) and proliferation was assessed in an AlamarBlue assay (n = 3). **(B)** A673 cells were treated with HPI-1 [20 µM], CK1D008 [0.3 µM, 1 µM] or SR-3029 [0.1 µM, 0.3 µM] and proliferation was assessed in an AlamarBlue assay. **(C)** PANC1, A549, LNT-229, HT29, HCT15 and HCT116 cells were pre-treated with CK1D008 [0.1 µM, 0.3 µM, 1 µM] for 48h, seeded in limited dilutions and cultured under anchorage-dependent non-proliferating conditions. Colony number was assessed after 9-11 days (n = 3). **(D)** Representative Western blot analysis of GLI1 protein levels in A673 cells lentivirally transduced with sgGLI1 or sgCtrl. **(E)** Knockout of GLI1 reduces anchorage-independent growth of A673 cells in soft agar. Representative images of formed anchorage-independent 3D tumorspheres (left panel), number of colonies (right panel) (n = 5). **(F)** A673 cells were lentivirally transduced with sgGLI1 or sgCtrl and proliferation was assessed in an AlamarBlue assay. **(G)** Representative Western blot analysis of overexpressed HA-tagged GLI1 and HA-tagged GLI2act protein levels in A673 cells. **(H)** PANC1 cells were pre-treated with CK1D008 [0.3 µM] for 48h. $1x10^6$ viable Ctrl or CK1D008 pre-treated PANC1 cells were engrafted in nude mice and tumor volume was monitored (n = 10). Student's t test was used for statistical analysis (*P < 0.035; **P < 0.002; ***P < 0.0002).
