## Supplementary material for "Casein kinase 1D encodes a novel drug target in Hedgehog-GLI driven cancers and tumor-initiating cells resistant to SMO inhibition": suppl tables

Supplementary Tables

**Table S1: Nucleotide sequences of primers**

| Target | Forward Primer (5'-3') | Reverse Primer (5'-3') |
|---|---|---|
| RPLP0 | GGCACCATTGAAATCCTGAGTGATGTG | TTGCGGACACCCTCCAGGAAGC |
| GLI1 | TCTGGACATACCCCACCTCCCTCTG | ACTGCAGCTCCCCCAATTTTTCTGG |
| PTCH1 | TCCTCGTGTGCGCTGTCTTCCTTC | CGTCAGAAAGGCCAAAGCAACGTGA |
| HHIP | ACTTGCCGAGGCCATATTCCAGGTT | ATCCCCACTATGCAGGGCACCAAC |
| CSNK1D | TTTCTGCCGTTCCTTGCGTTTTGAC | GTGTGAGGTAGGGGTGAGGGGTGTG |

**Table S2: Western Blot antibodies**

| Target (clone) | Supplier |
|---|---|
| anti-GLI1 (V812) | Cell Signaling, Danvers, MA, United States |
| anti-p44/42 MAPK (Erk1/2; tERK) | Cell Signaling, Danvers, MA, United States |
| anti-HA-Tag (C29F4) | Cell Signaling, Danvers, MA, United States |
| anti-β-Tubulin (9F3) | Cell Signaling, Danvers, MA, United States |
| anti-CSNK1D (AF12G4) | abcam, Cambridge, United Kingdom |
| anti-GLI2 (H-300) | Santa Cruz, Dellas, TX, United States |
| anti-rabbit IgG, HRP-linked | Cell Signaling, Danvers, MA, United States |
| anti-mouse IgG, HRP-linked | Cell Signaling, Danvers, MA, United States |
| anti-goat IgG, HRP-linked | Santa Cruz, Dellas, TX, United States |

**Table S3: Nucleotide sequences of single guide RNAs**

| Target | Targeting Sequence |
|---|---|
| GLI1 | AACTCGCGATGCACATCTCC |
| Non targeting control | GAGCTGGACGGCGACGTAAA |
